# Epigenetic switching outcompetes genetic mutations during adaptation to fluctuating environments

**DOI:** 10.1101/2021.03.11.434930

**Authors:** Dragan Stajic, Claudia Bank, Isabel Gordo

## Abstract

Epigenetic inheritance allows for the emergence of phenotypic plasticity in clonal populations and enables the rapid stochastic switching between distinct phenotypes. In natural environments, where stress conditions can recurrently fluctuate, clones with an epigenetic control of genes targeted by selection should be fitter than clones that rely solely on genetic mutation. To test this prediction, we engineered switcher and non-switcher yeast strains, where the uracil biosynthesis gene *URA3* is under fluctuating selection. Competitions of clones with an epigenetically controlled *URA3* with clones without switching ability (*SIR3* knock-out), show that epigenetic switching dominates under rapidly changing stresses. We further show that this advantage depends both on the switching rate and the period of environmental fluctuations. Remarkably, epigenetic clones with a high, but not with a low, rate of switching can co-exist with non-switchers even under a constant selective pressure, consistent with different constraints on the evolution of the rate of epigenetic switching.

## Introduction

Epigenetic inheritance comprises many interconnected molecular mechanisms that maintain development of distinct phenotypes through the regulation of gene transcription, independently of the underlying DNA sequence (Bird, 2007; Jablonka and Raz, 2009; Moazed, 2011). Epigenetically determined phenotypic states are typically less stable compared to those caused by genetic changes. The rate of epigenetic switching was shown to be several orders of magnitude higher than the mutation rate (Dodson and Rine, 2015; Van Der Graaf et al., 2015). Nevertheless, these states can be maintained for several generations before switching to the alternative phenotype (Johannes et al., 2009; Klosin et al., 2017).

Due to the transient nature of epigenetic inheritance, its potential contribution to the evolutionary processes is contentious (Charlesworth et al., 2017). However, theoretical and empirical studies have shown that the importance of the epigenetic switching during the process of adaptation is environment-dependent (Kussell and Leibler, 2005; Lachmann and Jablonka, 1996).

In a constant environment, where selective pressure is uniform during the course of adaptation, epigenetic control of gene expression acts primarily as a buffering mechanism until a more stable form of adaptation, i.e. genetic change, is acquired (Lachmann and Jablonka, 1996; Stajic et al., 2019; Torres-Garcia et al., 2020). Under such conditions, an epigenetic system of inheritance can influence adaptation (Kronholm et al., 2017) by maintaining a large enough mutational supply for beneficial mutations, consequently increasing the rate of adaptation (Klironomos et al., 2013; Kronholm and Collins, 2015). Epigenetic systems were also shown to change the spectrum of beneficial mutations during adaptation, enabling acquisition of adaptive mutations that modulate epigenetic control of gene expression (Stajic et al., 2019).

On the other hand, in fluctuating environments, cellular processes that induce and maintain phenotypic heterogeneity might have a predominant role during adaptation and provide a better long-term adaptive advantage than genetic mutations (Meyers and Bull, 2002). Here, epigenetically induced phenotypic heterogeneity could provide the basis for a bet-hedging survival strategy (Beaumont et al., 2009; Cohen, 1966), whereby alternative phenotypes are produced that might be maladaptive in one environment, but provide a net fitness benefit upon environmental change.

Indeed, theoretical work has shown that epigenetic switching should dominate over genetic changes in fluctuating environments, whereas genetic mutations are favored in more stable environments (Kussell and Leibler, 2005; Lachmann and Jablonka, 1996; Thattai and Van Oudenaarden, 2004). If the environment switches between two states periodically, the fitness advantage of a given binary epigenetic switching mechanism will depend on the rate of change between the two phenotypic states and the relative fitness difference of the two phenotypic states in the two environments. The optimal epigenetic switching rate is expected to be the one that matches the frequency of environmental fluctuations, *i.e.* 1/T, with T being the time interval of environmental change (Lachmann and Jablonka, 1996). As the environmental period becomes shorter, the optimal epigenetic switching rate increases. Under such conditions, genetic mutations between the two phenotypic states are expected to be of little effect due to their low probability of occurrence compared to the frequency of environmental oscillations (*e.g.* the genetic mutation rate in budding yeast tends to be several orders of magnitude lower than the epigenetic switching rate (Dodson and Rine, 2015; Lang and Murray, 2008)).

Recent experimental studies have shown that epigenetic switching can affect to certain extent the growth rate in fluctuating environments (Acar et al., 2008; Kronholm and Ketola, 2018; Proulx et al., 2019). Populations with a high rate of epigenetic switching have a higher growth rate upon an environmental change compared to populations with a lower rate of switching (Acar et al., 2008).

However, the direct measurement of the adaptive advantage of an epigenetic system of inheritance compared to genetic mutations, under different environmental conditions, has been difficult.

We designed an experimental setup in which theoretical predictions can be directly evaluated by competing a yeast strain containing an epigenetic machinery (with different rates of epigenetic switching) with a strain that can only adapt through genetic mutation (created via a knock-out of epigenetic silencing components). We ask two main questions: a) Under periodic environmental changes, can a strain capable of epigenetic phenotypic switching dominate over a strain that cannot switch?; b) How does the rate of epigenetic switching influence the adaptation dynamics in fluctuating environments?

We used previously constructed *Saccharmyces cerevisiae* strains in which a *URA3* reporter gene was inserted into a subtelomeric region (Stajic et al., 2019), resulting in differential epigenetic silencing of the gene. *URA3* is a widely used reporter gene that enables dual selection (i.e. selection for gene activation and inactivation). The gene is crucial for the production of uracil, which is essential for cell growth. However, in the presence of a drug, 5-Fluoroorotic acid (5-FOA), the activity of the Ura3 protein is deleterious, since it converts the drug into a toxic 5-Fluorouracil that kills the cell (Boeke et al., 1987). The strength of the silencing and the rate of switching between ON and OFF state of *URA3* expression in the subtelomeric region is dependent on the activity of Silent Information Regulator (SIR) proteins (Aparicio et al., 1991; Ivy et al., 1986; Rine and Herskowitz, 1987), which act as chromatin modifiers (Imai et al., 2000), and the relative distance of the gene from the telomere (Pryde and Louis, 1999). Using this well established system we selected for *URA3* gene activation (ON state) by removing uracil from the medium, or inactivation (OFF state) by adding 5-FOA.

We find that an epigenetic form of inheritance dominates over genetic mutations when yeast is propagated in fluctuating environments. However, this advantage of the epigenetic machinery is dependent on the switching rate. Surprisingly, we find that clones capable of epigenetic switching can coexist with non-switchers even in stable environmental conditions.

## Results

### An epigenetic system of inheritance dominates over mutations during adaptation to fluctuating environments

We chose a strain, referred to as fast epigenetic switcher, with subtelomeric *URA3* position that showed high levels of epigenetic switching between ON and OFF state: ON rate≈10^−2^, OFF rate≈10^−2^ (Stajic et al., 2019). We directly competed this strain with its corresponding Δsir3 mutant (referred to as non-switcher) that lacks an essential component of the SIR machinery and, consequently, lost its ability to epigenetically control gene expression in the subtelomeric region. Moreover, the non-switcher strain was previously shown to have a higher mutation rate than the wild - type (Stajic et al., 2019), which increases the probability of acquisition of beneficial mutations. This system allowed us to monitor, in a controlled manner, the effect of epigenetic changes and mutations during adaptation by determining the relative frequency of each strain in the population. To distinguish the two strains, each was marked with a different fluorescent marker; the epigenetic switcher with Red Fluorescent Protein (RFP) and the non-switcher strain with Yellow Fluorescent Protein (YFP). We first preselected the cells to be in ON state at the onset of the experiment, by growing the cultures in complete synthetic media (CSM) lacking uracil (fig. 1). We mixed the epigenetic switcher strain with the non-switcher in a proportion of 1:100 to mimic the case of invasion of an epigenetic strain into a non-epigenetic resident population and to minimize the probability of mutations in the epigenetic switcher background. To test the theoretical predictions, we exposed such co-cultures to fluctuating environments with periodicities equal or higher than the epigenetic switching rate, with alternating selection pressures for ON or OFF state of the *URA3* gene. Additionally, we also followed co-cultures in the two non-fluctuating environments with constant selection regimes. Theoretical studies predict the epigenetic switcher strain to dominate the populations in the fluctuating environments, especially in the environment in which the fluctuation frequency corresponds to the switching rate. In contrast, in the constant environments, in which a more stable form of inheritance would be favored, we expect populations to be dominated by the non-switcher strain.

**Fig. 1.**
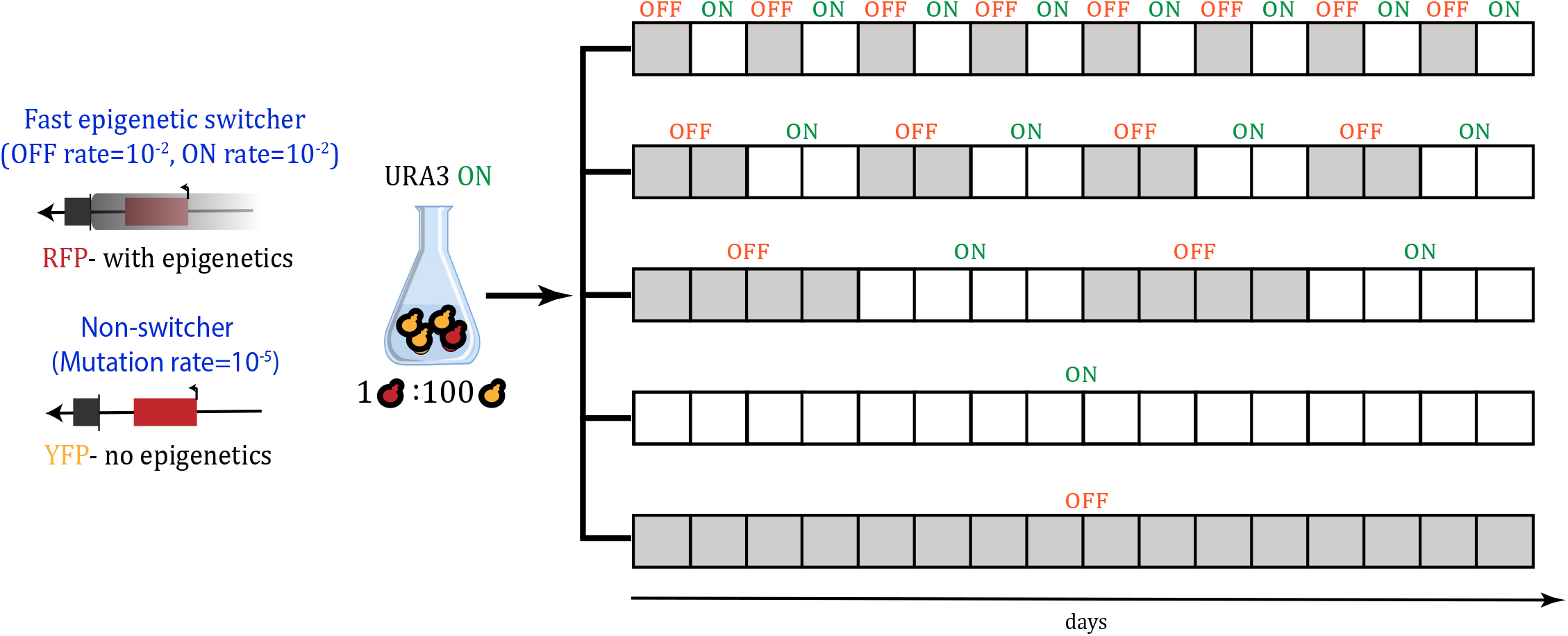
Experimental setup. Scheme showing experimental evolution setup used in the study. Yeast strains with epigenetic silencing (labelled with RFP) and a *SIR3* knock-out strain (labelled with YFP) were preselected in media without uracil (i.e. selection for active *URA3* gene) and mixed in 1:100 proportions, respectively. Subsequently, the mixed RFP/YFP yeast cultures were exposed to environments that fluctuated with different periodicity. Each box represents a 24h period after which populations were sampled and relative ratio of strains were determined. The color of the boxes represents the selection regime whereby grey boxes indicate period of selection for the inactive form of *URA3* and white colored boxes selection period for the active form of *URA3.*

In fluctuating environments, all of the 24 replicate populations survived during the course of the experiment, irrespective of the periodicity of environmental changes (fig. 2A). However, in fluctuating environments with 2 and 4 days periodicity, we observed larger oscillations in the total population size than in the fast, 1-day, fluctuating environment, especially during the periods that selected for the OFF state of *URA3* gene expression. Measurement of the relative frequency of the two strains within populations showed that despite the initial predominance of the non-switcher in the populations, after 96h the majority of cells were epigenetic switchers in all 24 populations of the three fluctuating environments (fig. 2B). However, after the early sweep of epigenetic switchers through the population, in some replicates the non-switcher recovered by the end of the experiment and increased in frequency above 50% (1 replicate in the 2-day fluctuating environment and 1 replicate in the 4-day fluctuating environment). This indicates that a genetic solution to the adaptation to fluctuating environments exists. Nevertheless, at the final time point of the experiment, the frequency of the epigenetic switcher was above 90% in the majority of populations exposed to fluctuating environments (23/24 in 1-day fluctuating environments, 23/24 in 2-day fluctuating environments, 20/24 in 4-day fluctuating environments). In contrast, under constant selection regimes only 4/24 populations in 5-FOA environment (selection for OFF state of URA 3) and 0/24 populations in the environment lacking uracil showed dominance of the epigenetic switcher after 96h hours (fig. 3B). To further quantify this, we compared the mean frequency of epigenetic switchers across all replicate populations for each environmental condition (fig. 4). The mean frequency of epigenetic switchers within populations was significantly higher in fluctuating environments as compared to stable environmental conditions. These results support the theoretical predictions that an epigenetic system of inheritance will have a strong beneficial effect in fluctuating environments, whereas genetic mutations are favored in stable environments.

**Fig. 2.**
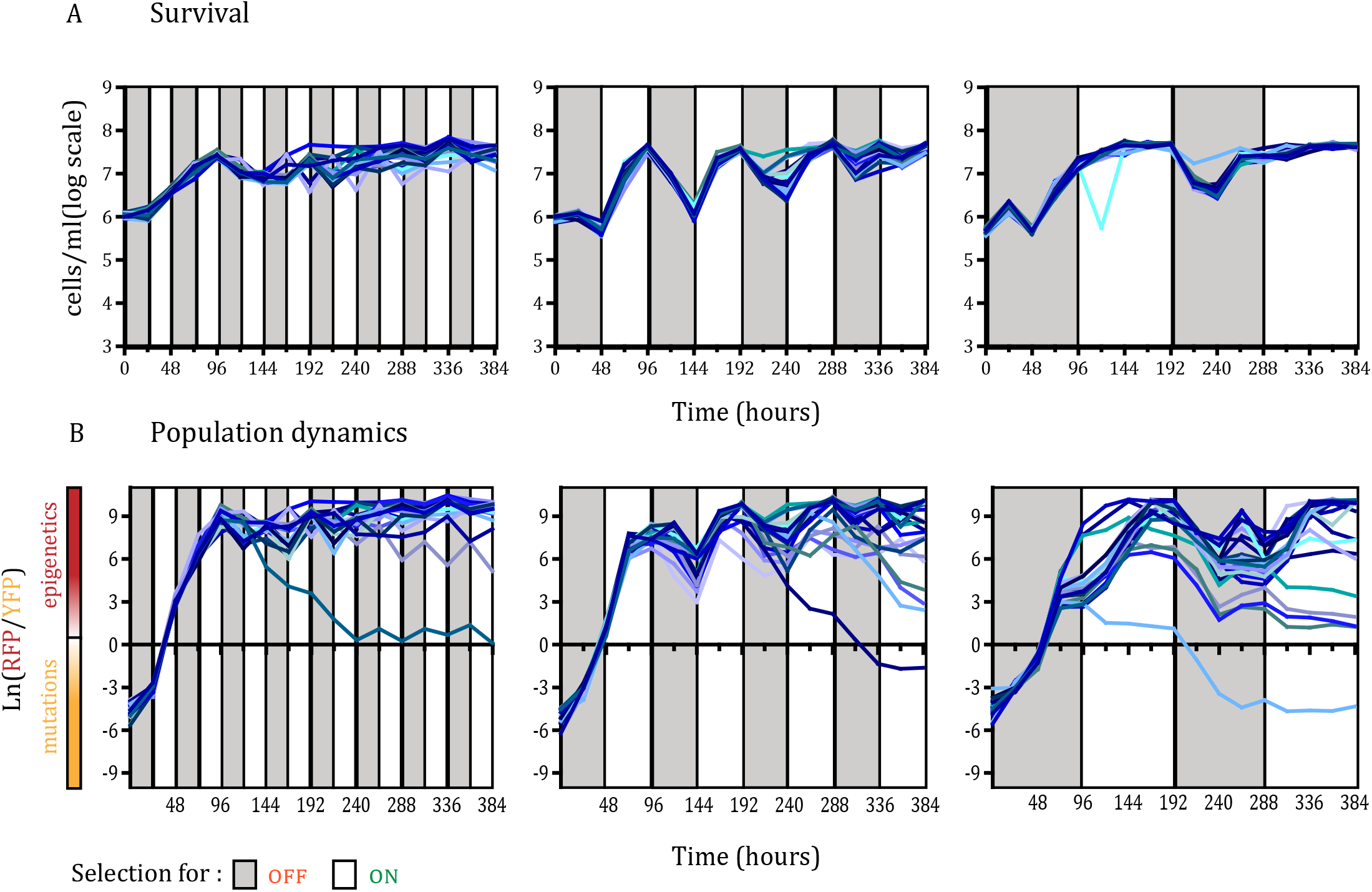
Epigenetic switching dominates over mutations in fluctuating environments. (**A**) Survival through the course of selection in the three fluctuating environments with distinct periodicities, determined using FACS methodology. Each line represents the number of cells in each replicate population (24 replicate populations for each environmental condition). Colored areas indicate the selection regime, grey corresponds to selection for inactive *URA3* and white for selection for the active form of the gene. (**B**) **D**ynamics of RFP/YFP ratios (with high rate of epigenetic switching) in fluctuating environments. The logarithm of RFP/YFP ratios for each of the replicate populations is shown, determined using FACS methodology. The color of the line for each population corresponds to the color of the lines in the survival graphs. Colored areas indicate the selection regime, grey corresponds to selection for inactive *URA3* and white to selection for active form of the gene. Positive values indicate dominance of the RFP strain, and negative values indicate dominance of the YFP strain.

**Fig. 3.**
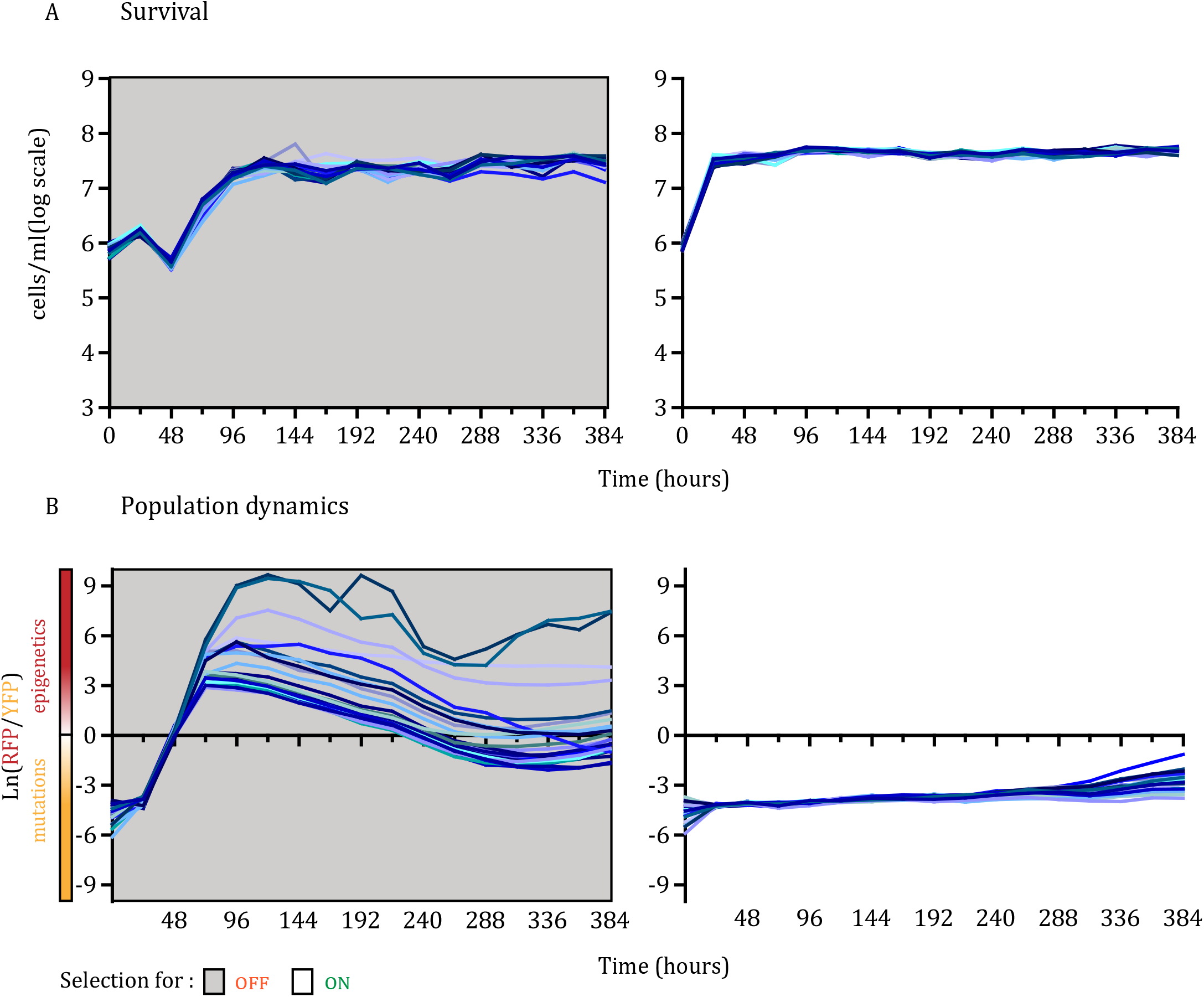
Epigenetic switching co-exists with mutations in stable environments. (**A**) Survival through the course of selection for two constant environments with distinct selection regimes here marked with the same colors as in the fluctuating environments (grey indicates selection for an inactive *URA3* gene and white for an active form of the gene), determined using FACS methodology. Each line represents the number of cells in each replicate population (24 replicate populations for each environmental condition). (**B**) Dynamics of RFP/YFP ratios in the two constant environments. The logarithm of RFP/YFP ratios for each of the replicate populations is shown, determined using FACS methodology. The color of the line for each population corresponds to the color of the lines in survival graphs. Colored areas indicate the selection regime as in panel A.

**Fig. 4.**
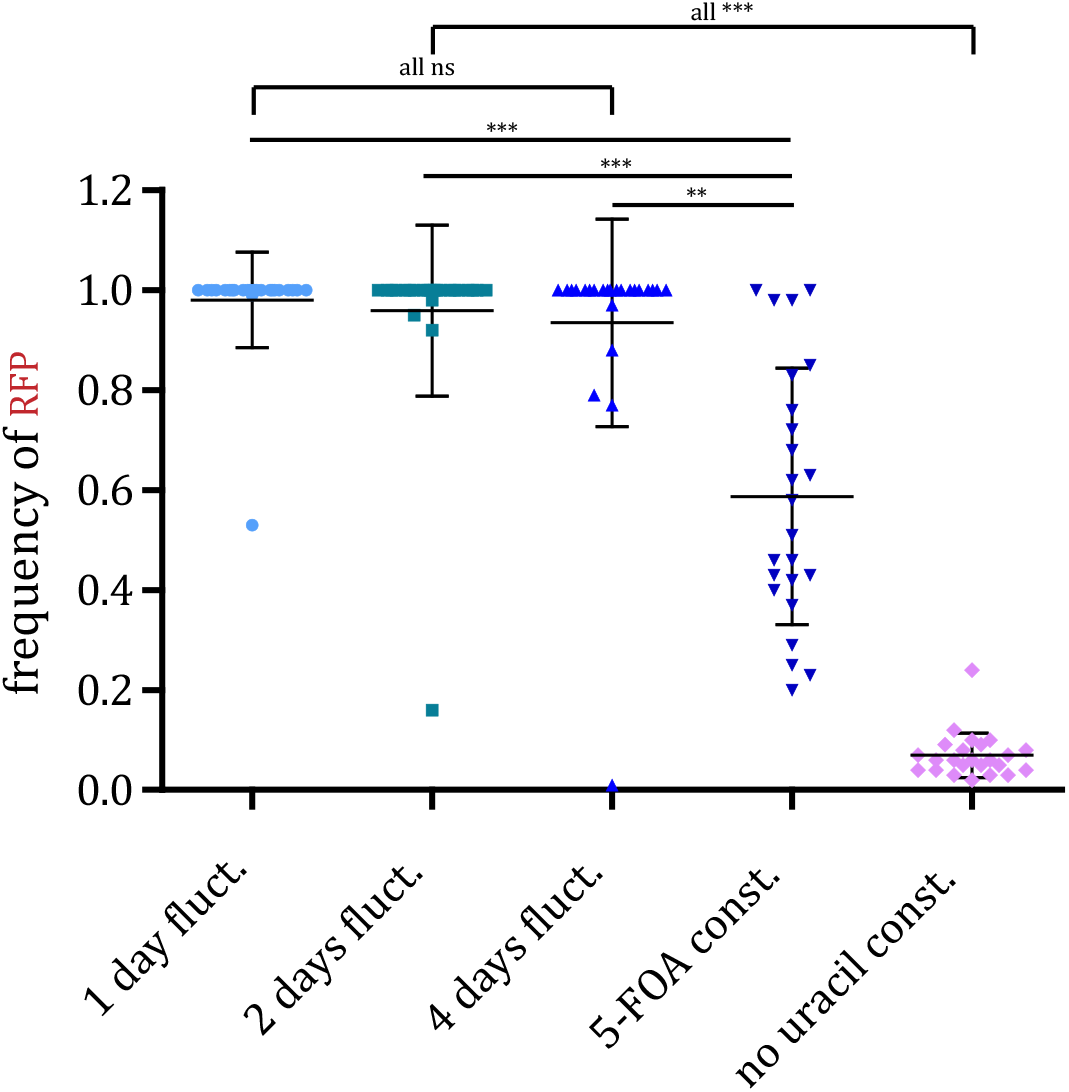
Populations selected in fluctuating environments show a higher frequency of the epigenetic strain than those grown in stable environments. We compared the frequency of the fast epigenetic switcher strain at the final time point across all selection regimes. Bars represent mean and standard deviation. Points represent frequenciess for each replicate population. The mean values between the selection regimes were compared using Dunn’s non-parametric test with Bonferroni correction for multiple testing.

Even though all of the replicate populations survived in the two constant selection regimes (fig. 3A), the population dynamics differed between them (fig. 3B). In the stable environment, in which we selected for the OFF state of *URA3* expression, we observed an increase in the relative frequency of epigenetic switchers in the populations in the first 72 hours (average frequency at 72h is 98%) due to the high rate of turning off *URA3* expression, which is consistent with the results in fluctuating environments. However, in stable environments this increase in frequency was followed by the appearance of beneficial mutations in the non-switcher background that decreased the relative proportion of epigenetic switchers (average frequency of epigenetic switchers at 384h is 59%). This indicates that despite the initial beneficial effect of epigenetic switching, in the long term epigenetically controlled gene expression states are hard to sweep to fixation, probably due to the cost of constantly switching to the less fit phenotype. Nevertheless, the epigenetic switcher persisted in the populations and was maintained at different frequencies until the end of the experiment (~200 generations; 384 hours), resulting in persistent phenotypic and genotypic heterogeneity in the populations.

### The adaptive importance of epigenetic machinery is environment-dependent

To confirm that the differences observed in the adaptive dynamics under the different environments are indeed due to epigenetic gene expression regulation, we determined the phenotypes of the evolved clones within each replicate population. Plating the replicate populations from the last time point on the non-selective (rich media) plates that were subsequently replica plated onto CSM media plates containing 5-FOA drug and CSM plates that lack uracil allowed us to phenotypically distinguish switchers from non-switchers. The growth of the evolved clones on both media indicates their ability to epigenetically switch between the active and inactive form of *URA3* gene. On the other hand, the inability of clones to grow on media lacking uracil would indicate a genetic inactivation of the uracil biosynthesis pathway. As our analysis of relative switcher/non-switcher frequencies indicated, epigenetic switchers were more common in fluctuating environments, whereas in stable environments genetic solutions were more prevalent (fig. 5A). To further confirm that the ability to switch between the two phenotypes indeed depended on the epigenetic machinery, we additionally replica plated the 5-FOA resistant clones onto media containing nicotinamide (NAM), a known inhibitor of SIR-mediated gene silencing (Bitterman et al., 2002). As expected, in fluctuating environments, especially under the 4-day fluctuation period, the periodicity of which corresponded to the switching rate, resistance to the drug is abrogated upon exposure to NAM. This shows that adaptation is dependent on the epigenetic machinery in these clones. On the other hand, drug resistance of the evolved clones from the stable environment is unaffected by the presence of NAM, providing further evidence of the genetic basis of adaptation to these environmental conditions (fig. 5B). Surprisingly, in several populations we observed clones that still showed the ability to switch between the two gene expression states, but in which resistance to 5-FOA was not altered by the addition of NAM. This indicates the existence of alternative adaptation mechanisms, independent from the epigenetic machinery, that maintained the switching ability without altering the uracil biosynthesis pathway directly.

**Fig. 5.**
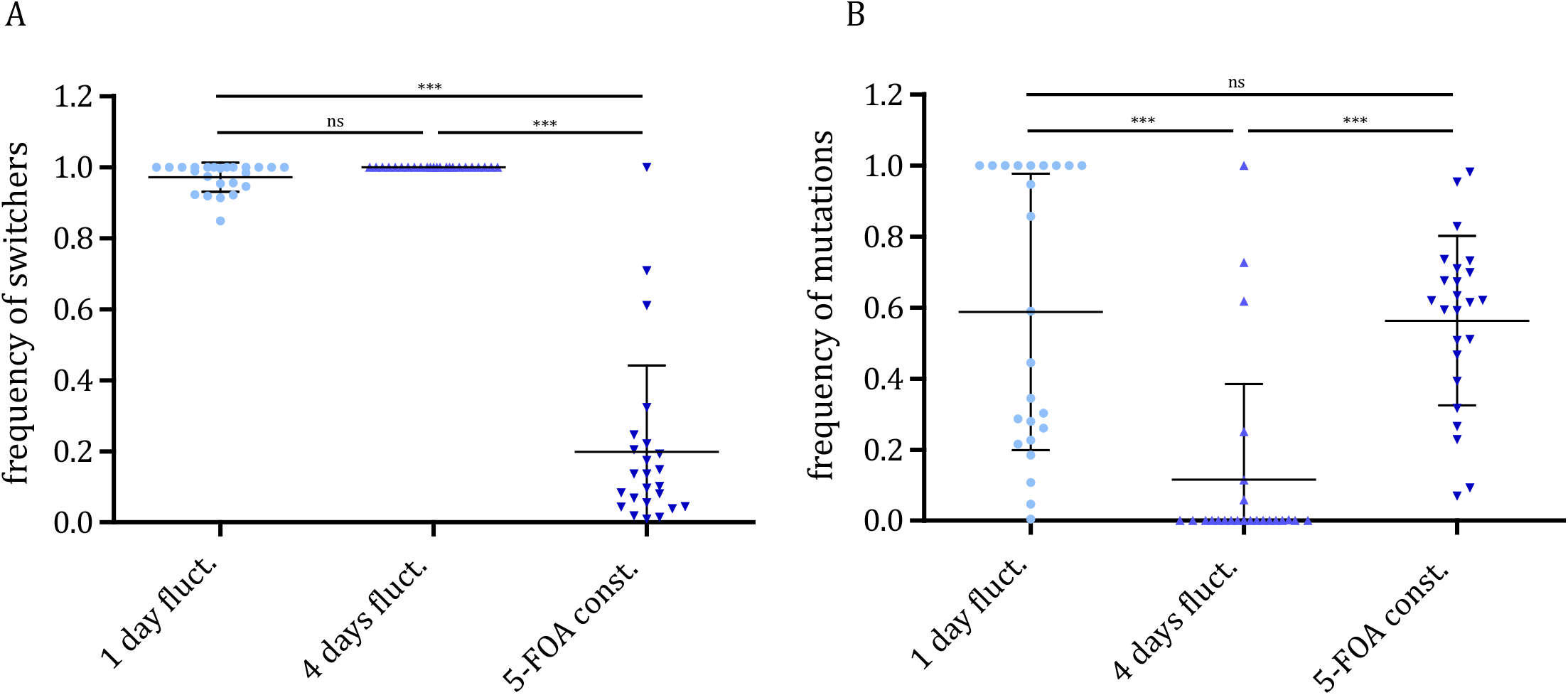
The adaptive advantage of epigenetic switching is dependent on the period of environmental fluctuation. (**A**) Points represent frequencies of clones that were able to grow on both the plates containing 5-FOA (selecting for OFF state of *URA3* gene) and the plates lacking uracil (selecting for the ON state of *URA3* gene) within each replicate population at the end of the experiment. Bars represent the mean and standard deviation. The mean values between the selection regimes were compared using Dunn’s non-parametric test with Bonferroni correction for multiple testing. (**B**) Points represent frequencies of clones whose 5-FOA resistance was not abrogated upon the addition of the inhibitor of epigenetic silencing, nicotinamide, within each replicate population. Bars represent the mean and standard deviation. The mean values between the selection regimes were compared using Dunn’s non-parametric test with Bonferroni correction for multiple testing.

### The effect of the epigenetic inheritance system depends on its rate of switching

To make sure that the effect of epigenetic switching observed in the fluctuating environments was solely due to the intrinsic characteristic of the epigenetic system of gene expression control and not due to possible deleterious effects of the *SIR3* knock-out mutation, we performed an additional evolution experiment. We competed a yeast strain, referred to as slow epigenetic switcher, that has a lower rate of epigenetic switching (ON rate≈10^−2^, OFF rate≈10^−6^) (Stajic et al., 2019), but otherwise has the same growth and mutation rate as the fast epigenetic switcher, with its corresponding Δsir3 mutant. The slow epigenetic switcher is preferentially in ON state of *URA3* expression and behaves very similarly to the non-switcher strain. Similar to the procedure in the first experiment, we preselected the cultures of the two strains in media lacking uracil and then exposed the co-cultures to the three fluctuating environments (fig. 1).

We observed that populations tended to go extinct as the environmental period increased (fig. 6). Under 1-day fluctuating periods, all populations survived, likely because the rapid environmental changes (every 10 generations) weakened the selective pressure, since cells spent less time in each environment. On the other hand, under 2-day fluctuations we observed extinction in 2/24 replicate populations, and under 4-day fluctuations all 24/24 replicate populations went extinct. This is a very different outcome than that observed in the co-cultures with the fast epigenetic switcher, for which under 4-day fluctuations all populations survived (P<0.00001, Fisher’s exact test). Thus, the rate of epigenetic switching plays an important role for the adaptation to fluctuating environments, further confirming theoretical predictions (Lachmann and Jablonka, 1996). If the epigenetic switching rate is low, the populations may go extinct because they cannot respond quickly enough to the strong selection pressure, whereas under fast epigenetic switching a more phenotypically heterogeneous population can be rapidly established to cope with fast environmental changes. Furthermore, we observed that the populations which survived the experiment in rapidly fluctuating environmental conditions were dominated by non-switcher strains. Indeed, in 21/24 surviving replicate populations under 1-day fluctuations and 19/22 surviving replicate populations under 2-day fluctuations showed a frequency of non-switcher strains that was above 90%. This is in contrast to the experiment with the fast switcher in which the majority of populations were ultimately dominated by the switcher strain, further demonstrating the power of an epigenetic system during adaptation to fluctuating environments.

**Fig. 6.**
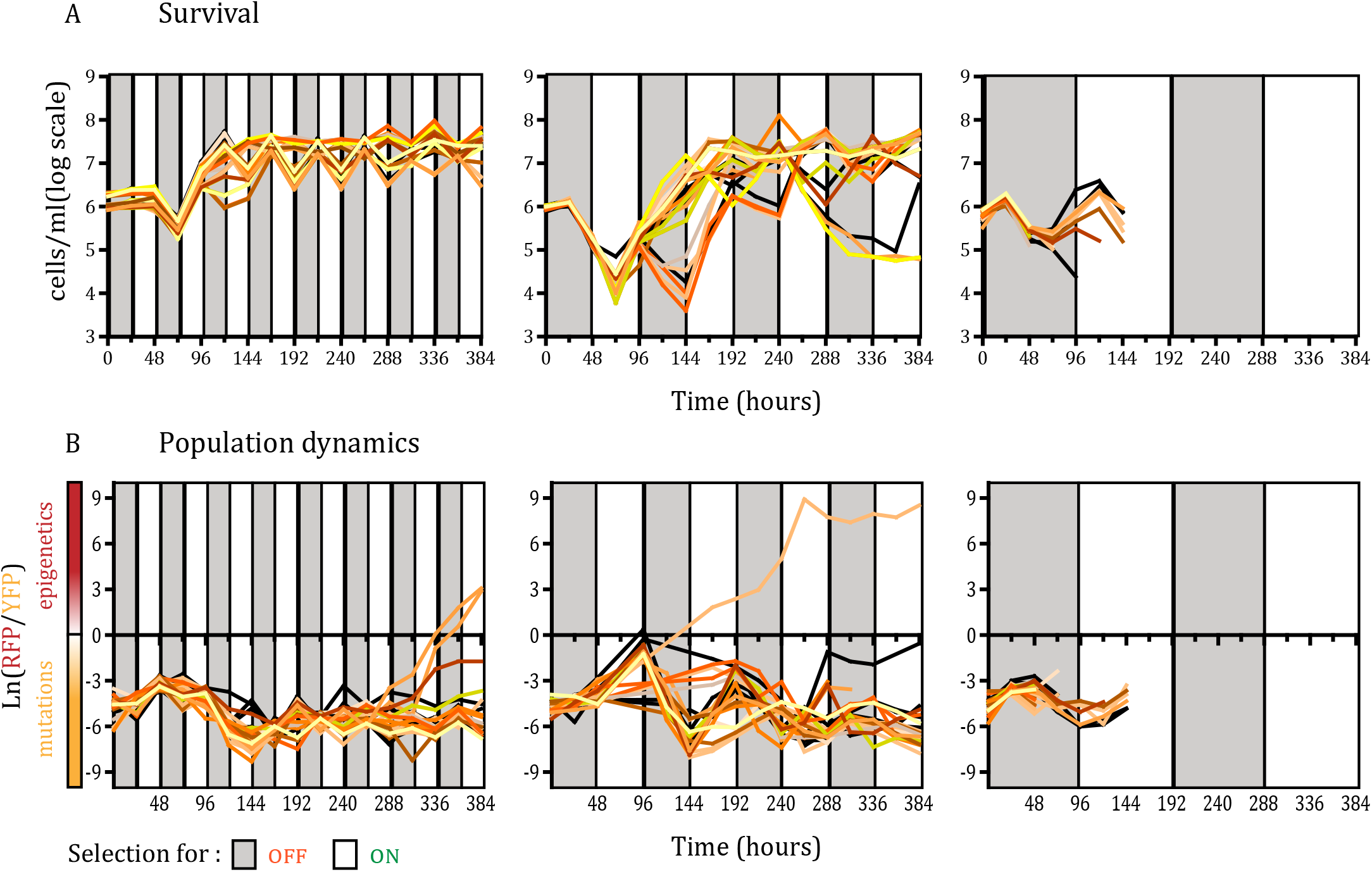
The dominance of epigenetic switchers depends on the rate of epigenetic switching. (**A**) Survival through the course of selection for a strain with a low rate of epigenetic switching in the three fluctuating environments with distinct periodicities, determined using FACS methodology. Each line represents number of cells in each replicate population (24 replicate populations for each environmental condition). Colored areas indicate the selection regime, grey corresponds to selection for inactive *URA3* and white for selection for the active form of the gene. (**B**) Dynamics of RFP/YFP ratios (with a low rate of epigenetic switching) in the fluctuating environments. The logarithm of RFP/YFP ratios for each of the replicate populations is shown, determined using FACS methodology. The color of the line for each population corresponds to the color of the lines in survival graphs Colored areas indicate the selection regime as in panel A. Positive values indicate dominance of RFP strain, and negative dominance of YFP strain.

## Discussion

Epigenetic control of gene expression can have profound effects on phenotypic variation (Jablonka and Raz, 2009) and provide a bet-hedging survival strategy in a novel environment. Due to the fast rates of switching between different selectable phenotypes, an epigenetic system of inheritance effectively establishes phenotypic heterogeneity in the population, whereby it produces phenotypes that might be maladaptive in one environment but beneficial upon an environmental change (Bódi et al., 2017). In stable environments, where populations are under constant uniform selective pressure, epigenetic systems offer a buffering mechanism until adaptive mutations are acquired. Indeed, our results support these findings. In a constant environment selecting for *URA3* inactivation (5-FOA stable environment), we observed an initial increase in frequency of epigenetic switcher (up to 98% at 72h) that was subsequently followed by the acquisition of mutations by the end of the experiment. Surprisingly, we observed that the epigenetic switcher strains were still maintained in the population under stable environmental conditions, even after 200 generations, thereby maintaining phenotypic and genetic diversity in the population.

Carriers of an epigenetic system of inheritance were found to increase in frequency in fluctuating environments. This was especially true when the rate of epigenetic switching corresponded to the frequency of the environmental changes. We observed that fast epigenetic switchers dominated in all fluctuating environments, regardless of the periodicity of fluctuations. However, the molecular basis of this phenotypic switch was different between fluctuation periods. For example, in the environment in which the fluctuation periodicity corresponded to the switching rate (4-day fluctuations), the switch was still dependent on the epigenetic machinery at the end of the evolution experiment, whereas in an environment with a faster fluctuation period (1-day fluctuations) the molecular basis of the switch seems to have been genetically altered.

Our results therefore indicate that epigenetic switching and the resulting phenotypic variation are environment-dependent. Ecological studies of epigenetic variation in nature seem to support this (Richards et al., 2017). Additionally, the correlation between particular epigenetic gene expression patterns and the habitat seems to be species-specific (Alonso et al., 2015; Niederhuth et al., 2016). Additionally, it remains difficult to draw a causal link in natural populations between adaptation and particular epigenetic marks (Herrera and Bazaga, 2010)

Furthermore, our results indicate that the strength of epigenetic gene activation or inactivation depends on the essentiality of the gene in a given environment. In other words, stress response genes that are essential for quick physiological response to a sudden environmental insult and that are otherwise non-essential under normal, stable conditions, are expected to be under epigenetic transcriptional regulation. That seems to be the case in budding yeast, where most of the genes in the subtelomeric region that were shown to be under epigenetic control are stress related genes (Ellahi et al., 2015).

In sum, our experimental set-up offers a controllable and tractable system by which we can monitor the effects of heritable gene expression states and mutations during adaptation to different environments. Our study provides direct evidence of the importance of an epigenetic system of gene expression during adaptation to fluctuating environments. This system might provide additional opportunity for populations to adapt to rapidly changing conditions and prove crucial for survival under adverse environmental fluctuations that are expected to increase with climate change. Additionally, epigenetic switching, as we have shown, can provide an additional layer for the maintenance of phenotypic and genotypic heterogeneity in the population, which represents the basis for evolution.

## Materials and Methods

### Yeast strains and growth conditions

All *S. ceverevisiae* strains used in this study were derived from the S288c background. The fast epigenetic switcher strain (YIG1) was constructed by the integration of a NatMX6(noursethricin resistance)-mCherry cassette into the original *URA3* locus (between 115,929 and 117,048 position on chromosome V) of the LJY186 strain (*MATα, trpΔ63*, *hisΔ200, ura3Δ::KanMX6, TEL-XIL::URA3 position 1373*), deleting the originally positioned Kanamycin resistance cassette (KanMX6). The NatMX6-mCherry cassette was amplified from pDS3, constructed by insertion of an mCherry gene from pLJ760 into pAG25, containing a NatR cassette. The slow epigenetic switcher strain (YIG2) was constructed similarly, by the integration of an NatMX6-mCherry construct into the original *URA3* locus of the LJY185 strain (*MATα, trpΔ63*, *hisΔ200, ura3Δ::KanMX6, TEL-XIL::URA3 position 1623*). The YIG1 corresponding non-switcher strain (YIG3) was constructed by the integration of a NatMX6-mCitrine cassette into the original *URA3* locus of the LJY193 strain (*MATα, trpΔ63*, *hisΔ200, ura3Δ::KanMX6, sir3Δ::HYG, TEL-XIL::URA3 position 1373*). The natMX6-mCherry cassette was amplified from pDS4, constructed by insertion of the mCitrine gene from pLJ761 into pAG25, containing a NatR cassette. The YIG2 corresponding non-switcher strain (YIG4) was constructed by the integration of a NatMX6-mCitrine cassette into the original *URA3* locus of the LJY192 strain (*MATα, trpΔ63*, *hisΔ200, ura3Δ::KanMX6, sir3Δ::HYG, TEL-XIL::URA3 position 1623*). The original plasmids and yeast strains used in the design and construction of strains in this study were a kind gift from Lars Jansen lab.

All strains were maintained either in rich medium [YPD; 1% Bacto yeast extract-BD (Fisher; #212720), 2% Peptone (Fisher; #BP1420-500), 2% Glucose (Merck; #1.08342.1000), either as liquid medium or supplemented with 2% Agar (Roth; #2266,4) for solid medium] or in complete synthetic dropout medium [CSM; 0.7% Yeast nitrogen base (Sigma; #Y0626), 0.1% Complete synthetic medium (MP Biomedicals; #4560-222), 0.005% Tryptophan (Sigma; #T0254), 0.002% Histidine (Sigma; #H8000), 2% Glucose (Merck; #1.08342.1000) as liquid medium].

### Experimental evolution

All strains were preselected to be in *URA3*^+^ state by growing cells in liquid CSM lacking uracil for 16 hours at 28°C. Next, around 10^6^ cells in a 1:100 proportion of epigenetic switcher and non-switcher strain were diluted into 1ml liquid CSM containing 5-FOA [CSM supplemented with 0.05% 5FOA (Apollo Scientific; #PC4054) and 0.001% Uracil (Sigma; #U0750)]. Each day, 100μl of the culture was placed into fresh media. Depending on the periodicity of environmental fluctuations (see fig. 1) media was alternated between CSM containing 5-FOA and regular CSM. At each time point, 10μl of the cultures was mixed into 190μl of 1%PBS solution containing SPHERO fluorescent spheres (AccuCount 2.0 μm blank particles) that enabled accurate determination of the volumes. This mix was subsequently analyzed using Flow Cytometry to determine the number YFP and RFP labelled cells in each population. Exact number of cells in each population was obtained by the multiplication of detected events with proper dilution factor. If the total population size was less than 10^5^ cells for more than 4 time points, the population was considered extinct.

### Flow Cytometry

Flow Cytometry was performed in a BD LSR Fortessa™ SORP flow cytometer, using a 96-well plate High-Throughput Sampler (HTS). The relative number of respective fluorescently labelled yeast cells in each replicate population was determined by the number of counts of detected fluorescent events and the appropriate dilution that was made in PBS solution. The instrument is equipped with 488nm laser for scatter parameters and YFP detection and 561nm laser for mCherry detection. Relative to the optical configuration, YFP and mCherry were measured using bandpass filters in the range of 540/30 nm and 630/75nm, respectively. The analyzer was also equipped with a forward scatter (FSC) detector in a photomultiplier tube (PMT) to detect yeast. The results of the measurements were analyzed using Flowing Software version 2.5.1, developed by Perttu Terho, University of Turku. All Flow Cytometry experiments were performed at the Flow Cytometry Facility of Instituto Gulbenkian de Ciência, Oeiras, Portugal. The data from Flow Cytometry analyses is available in Supplemental Table 1.

### Phenotypic characterization of the evolved clones

We plated appropriate dilutions of each replicate population from the last time point on the rich media plates. The dilutions were made using the total cell numbers determined by flow cytometry so that around 100 cells were plated. The plates were incubated at 28°C for 3 days. Subsequently, the colonies were counted and replica plated onto CSM plates containing 0.1% 5-FOA as well as onto regular CSM plates (without 5-FOA and without supplemented uracil). These plates were incubated at 28°C for 5 days, after which period the cells were counted. The frequency of epigenetic switchers was determined by dividing the number of colonies that grew on both CSM plates and the number of colonies on the rich media plates. Furthermore, the colonies from 5-FOA containing plates were replica plated on CSM plates that contained 0.1% 5-FOA and were additionally supplemented with 5mM nicotinamide (Fisher; # 1663C). These plates were incubated for additional 5 days at 28°C, after which the colony number was scored. The frequency of clones with genetic changes was determined by the division of the colony number from plates with nicotinamide and colony number from the original rich media plates.

## Acknowledgments

We would like to thank Lars Jansen (University of Oxford, UK) for the strains and helpful comments. We would also like to thank Davide Cusseddu and all members of Evolutionary Biology and Evolutionary Dynamics groups, as well as, Rike Stelkens (Stockholm University, Sweden) for their support and ideas. We acknowledge the Flow Cytometry Facility of the Instituto Gulbenkian de Ciência for their support. D.S. was supported by Fundaçao para a Ciência e a Tecnologia (FCT) PREPARE project (JPIAMR/0001/2016-ERA NET), to I.G. and C.B., by FCT Project PTDC/BIA-EVL/31528/2017 to I.G. and ONEIDA and Congento projects (LISBOA-01-0145-FEDER-016417 and LISBOA-01-0145-FEDER-022170), both co-funded by FEEI - “Fundos Europeus Estruturais e de Investimento” from “Programa Operacional Regional Lisboa 2020” and FCT. D.S. is also grateful for support from Wallenberg Foundation (project grant: 2017.0163). C.B. is grateful for support by EMBO Installation Grant IG4152 and by ERC Starting Grant 804569 - FIT2GO from the European Research Council.

## Data availability

The data from Flow Cytometry measurements are available in Supplemental Table 1. All the strains used in the study are available upon request.

## Author contributions

D.S., C.B. and I.G. conceived the study and designed the experiments. D.S. constructed the strains and performed the experiments. D.S., C.B and I.G. critically analyzed the data. D.S., C.B. and I.G. wrote the manuscript. C.B. and I.G. provided resources, funding and supervision.

